# *Plasmodium* killing and antimicrobial lineal or cyclic-peptides computationally targeted the circumsporozoite CSP protein

**DOI:** 10.1101/2025.09.30.679691

**Authors:** J Coll

## Abstract

The RT39 **a**mphipathic **l**ineal-**p**eptide (ALP), described as the faster membranolytic-killer of circumsporozoite *Plasmodium falciparum* (Aguirre-Botero *et al*, 2024), computationally predicted the highest affinities to the external **c**ircum**s**porozoite **p**rotein (CSP), compared to other described ALPs. Additional computational co-cyclizations of CSP/ALPs generated CSP/cycRT39 cyclic-conformers predicting the highest picoMolar affinities to target CSP-TSR domains with *de novo* alternative amino acid sequences, proteolytic resistance, and chemical stability. A preliminary co-screening of Uniprot antimicrobial peptides (AMPs), discovered some CSP/cycAMP-conformers predicting similar high affinities. These novel cycALP and/or cycAMP amino acid sequences may deserve experimental validation tests. Could some of them help to advance on circumsporozoite interference in mosquitos and/or humans?.

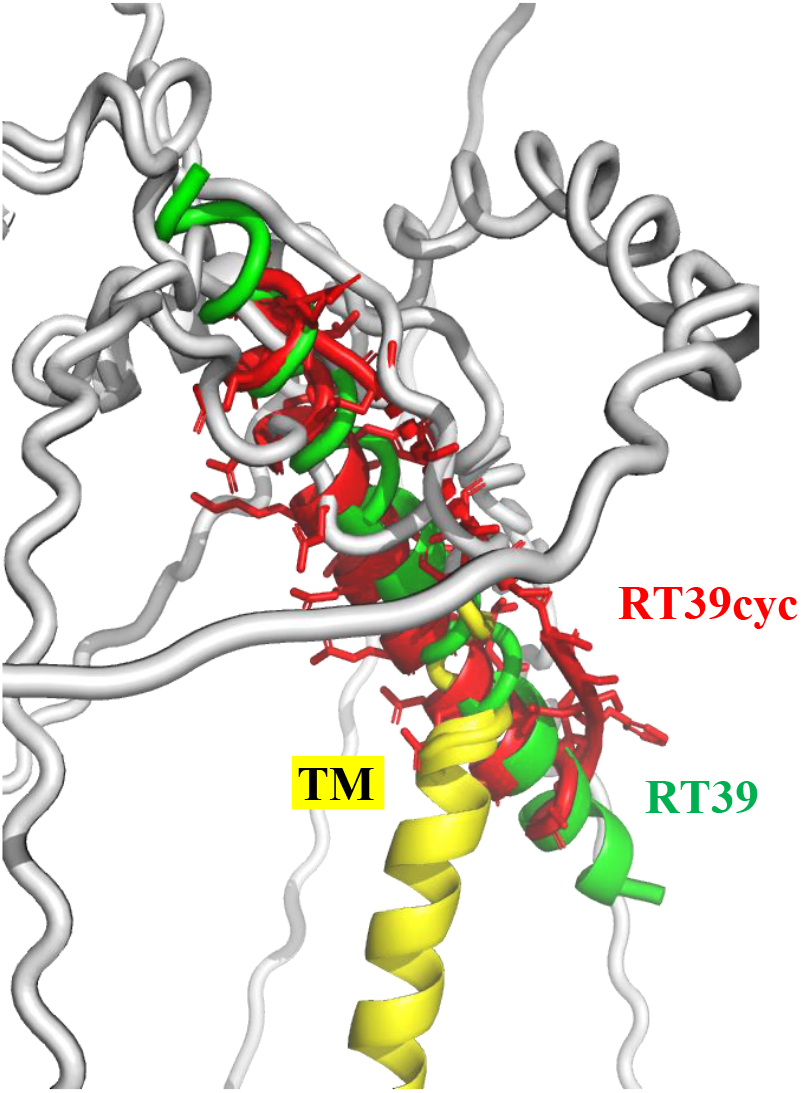

## *Plasmodium* CSP and amphipathic lineal-peptides (ALP)

Only 10-100 gliding-elongated cells of *Plasmodium falciparum* protozoan circumsporozoites (circulating sporozoites) are inoculated into the human skin per mosquito’s bite^1^. The inoculated circumsporozoites migrate through human blood to reach the liver, multiply to thousands of rounded merozoites per hepatocyte^1^ that infect and replicate into blood erythrocytes^2^ and finally cause clinical deathly infections^3^ . <u>**C**</u>ircum<u>**s**</u>porozoites are densely coated by a **p**rotein (CSP) which has been targeted by numerous prevention vaccines^10, 11^ and therapeutic antibodies and/or small drugs (including computational predictions of RFdiffusion peptides^4^ ).

Lab-designed <u>**a**</u>mphipathic <u>**l**</u>ineal-<u>**p**</u>eptides (ALP) have been recently optimized for fast *in vitro* membranolytic-killing of *Plasmodium* circumsporozoites (**Table 1**) ^5^. The designed ALPs belong to the <u>**A**</u>nti<u>**m**</u>icrobial <u>**p**</u>eptides (AMP) group of peptide structures. AMPs are membranolytic (membrane-permeabilization) defensive molecules widespread in uni- and multi-cellular organisms, contributing to killing many invading pathogens, such as pathogenic bacteria, fungi, parasites and/or viruses. Some AMPs are employed in medicine, food preservation, and/or agriculture.

**Table 1.**
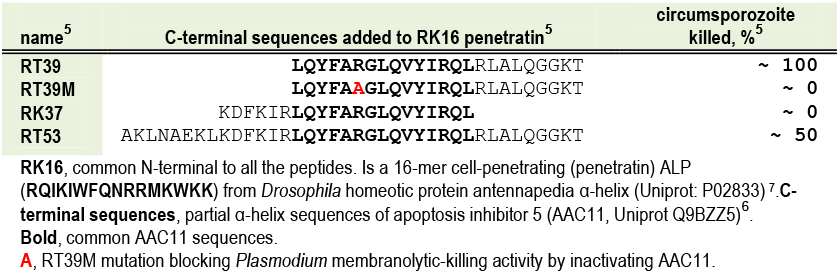
Amphipathic Lineal-Peptides (ALP) reported against circumsporozoites^5^.

The RT39 ALP was designed by fusing the penetratin cell-translocator (RK16) to selected heptad-repeat domains of the cell-penetrating AAC11 α-helix 18 peptide^6^. RT39 killed microsporozoites faster than anti-CSP antibodies^5^.

The RT39 activity extends from an unique circumsporozoite membrane binding/permeabilization event to induce gliding-immobilization and membranolytic-killing. In contrast, the RT39M mutant, substituting Arginine R^22^ by Alanine (RT39 R^22^A), blocked membranolytic-killing activity. Longer-peptide sequences such as RT53 were also membranolytic-killing circumsporozoites^6, 7^ but showed weaker and slower activity than RT39 (**Table 1**)^**5**^. High membranolytic activities may depend on several RT39 molecules inducing pores. In such case, the RT39M or RT53 lower pore-forming activities, could also be explained by inter-peptide reduced interactions^8^.

Among the many studies exploring the properties of anti-CSP antibodies induced by immunization or vaccines, only those with < nanoMolar affinities could inhibit malaria infections in mice^9-14^ and/or in most recent vaccines^15, 16^. Some of these antibodies that inhibit *P*.*falciparum in vitro* and in mice models^18, 19^, crystallographycally demonstrated that binding within picoMolar affinities that could change the conformations of the CSP NANP disordered-repeats^17, 18^ (7uym.pdb, 7v05.pdb, protein data bank structures). In contrast to anti-CSP, antibodies targeting TSR were rarely described, most probably because of their masking during migration ^20^. Therefore, all these results suggest that targeting CSP-TSR with the highest affinities, may increase the chances to interfere *in vitro* and/or *in vivo* with *Plasmodium* circumsporozoites.

Complementing present results with *Plasmodium* vaccines^10,11,19-21,31,32,33,22,37^and/or therapeutic anti-CSP antibodies^23^, CSP could also be targeted by small drug molecules, including peptides. Our previous attempts included *de novo* predictions of small drug-like compounds by computationally mimicking co-evolution^26^ to penetrate faster and deeper into the vast drug-chemical space^6, 7,8^, rather than screening large molecular banks^24, 25^. Such attempts targeted only the crystalyzed CSP-TSR model, following similar previous work on other protein-ligand pairs ^26-30^. Alternatively, computational *de novo* co-generated lineal and cyclic peptides predicting picoMolar affinities were also reported targeting either the crystallized CSP-TSR model ^26^ or full-length CSP Alphafold2 models^4^, respectively.

Cyclic-peptides predicting high affinities to CSP could be administered perhaps both to mosquito aquatic larvae (targeting their hemolymph and salivary glands) and/or to humans at high-risk of infection (targeting skin, blood, and liver). Targeting circumsporozoites would require the highest-affinity cyclic-peptides to: **i)** minimize side-effects, **ii)** deal with short-time exposures, **iii)** minimize proteolytic degradations, **iv)** maximize chemical stabilities, and/or **v)** minimize membrane permeability challenges by targeting external CSP^31-38^.

Although previous explanations for RT39 membranolytic-killing included mostly CSP-independent hypothesis^6^, this work explored for CSP-dependent docking of ALPs and AMPs. The computational strategy included :

1. Alphafold2 CSP/ALP co-modeling^39-41, 4^, ALPs from previous report^5^,
2. CSP/cycALP-conformer cyclic-ALPs co-generations,
3. Alphafold2 CSP/AMPs co-modeling, AMPs from Uniprot bank, and
4. CSP/cycAMP-conformer cyclic-AMPs co-generations.

## Computational results

### Amphipathic lineal-peptide (ALP) computational affinities and contacts to *Plasmodium* CSP

Five conformers (named from _1 to _5) of <u>**C**i</u>rcum<u>**s**</u>porozoite <u>**p**</u>rotein (CSP) complexes with each of the <u>**a**</u>mphipathic <u>**l**</u>ineal-<u>**p**</u>eptides (ALP): RT39, RT39M mutant, RT53 longer-peptide, RK37 shorter-peptide and RK16 penetratin (**Table 1**)^5^, were generated by Alphafold2. CSP/ALP-conformer energies, ∼ affinities and amino acid 4 Å contacts were predicted by Prodigy.

Results predicted an exceptionally docking-score of -27.4 Kcal/mol (maximal affinity) only for the CSP/RT39_3 conformer (**Figure 1, RT39_3 red**). In contrast, the other CSP/RT39_1, _2, _4, or _5 conformers (**Figure 1, violet**), and all the conformers from RT39M, RT53, RK37 or RK16 (**Figure 1, cyan, orange, gray**, and **green**), predicted docking-scores ranging between -7.2 to -16.7 Kcal/mol (lower affinities). The highest affinities predicted by 20 % of the RT39 conformers (one out of five) and the lower affinities of all other ALPs, correlates with their reported circumsporozoite highest membranolytic-killing activity and the limited location of its membranolytic beginnings^**5**^. These CSP/RT39_3 conformer highest affinities compared to those from other ALPs, may be explained by its unique targeting to all CSP-TSR residues and centered NANP-repeats (**Figure 1, red rectangles**), compared to other CSP/ALP conformers which targeted discontinued TSR stretches and none NANP repeats (except RT39M_5).

**Figure 1.**
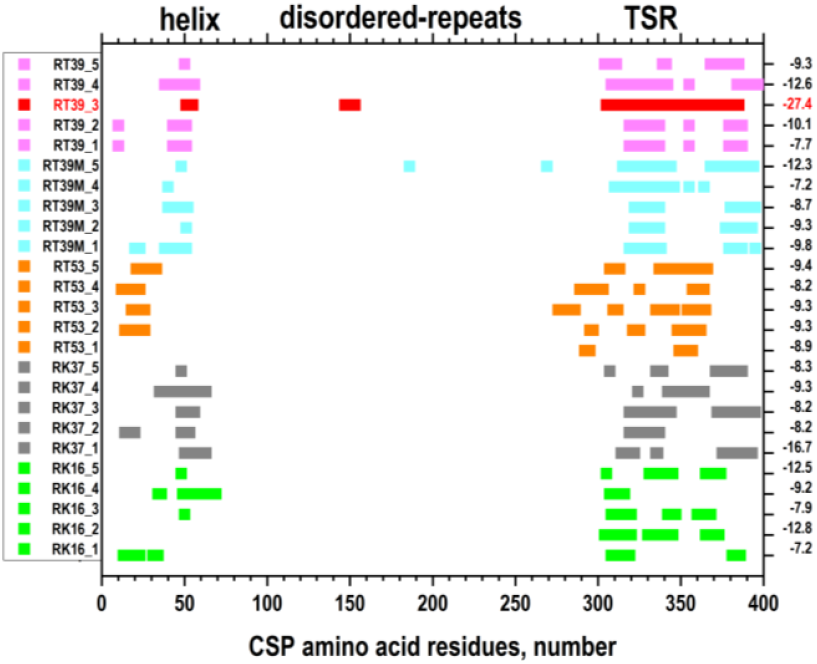
CSP amino acid contacts to ALPs and their affinities. CSP was Alphafold2 co-modeled with ALPs (**Table 1**) All conformer affinities were merged for the ranks, to generate five CSP/ALP conformers per ALP (named from _1 to _5). The CSP N-terminal residues predicted a signal peptide (SP) followed by an helix (45-55 residues)^42^, the hepatocyte surface binding HSPG (85-92) and the protease cleavage ^93^KLKQP (93-97)^43^ site. The NANP disordered repeats expanded ∼ half of the CSP (105-272). The C-terminal residues contained the crystallized cell-adhesive TSR (3vdj.pdb) (312-324)^8.^, the CS-flap (348-363)^24^, two highly conserved disulphide bonds (^338^C-^369^C and ^342^C-^374^C) and the transmembrane TM α-helix (376-396). **Left column**, Alphafold2 co-modeled CSP/ALPs conformers (_1 to _5). **Right column**, Prodigy-generated affinities in ∼ Kcal/mol. **RT39_3**, CSP/RT39 conformer predicting < picoMolar highest affinities (n=3).

The number of the CSP/RT39_3 conformer contacts were higher for its N-terminal penetratin (7-17 contacts for the 1-16 RT39 residues), than for its C-terminal AAC11 (1-11 contacts for the 17-32 RT39 residues and no contacts for the 33-30 RT39 residues)(**Figure 2**).

**Figure 2.**
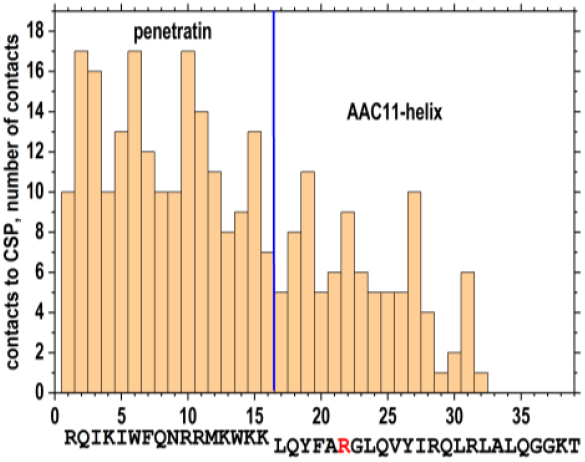
Number of RT39_3 amino acid contacts in the CSP/RT39_3 conformer. RT39 was designed by fusing penetratin (RK16) with AAC11 α-helices (**Table 1**). The CSP/RT39_3 conformer was generated by providing both CSP and RT39 amino acid sequences (n=3) to Alphafold2. Contacts were calculated by Prodigy. **Blue vertical line**, virtual limit between the two-fused peptide sequences in RT39. **R**, Arginine to Alanine mutation blocking circumsporozoite membranolytic-killing activity in RT39M.

### ALP cyclizations

To limit ALPs proteolysis and favour their chemical stability, six cyclic-conformers (CSP/cycALP) per CSP/ALP were generated. The new CSP/cycALP-conformers were labeled by adding _0 to _5 to each name (i.e. RT39_3_3). Most of the new CSP/cycALP-conformer predicted higher affinities up to ∼ 10^-5^ picoMolar (low binding-scores), compared to their CSP/ALP counterparts (**Figure 3, solid** *vs* **open red circles**), confirming previous reports^42, 51, 4^. The partial conservation of many of the CSP/ALP contacts in CSP/cycALP, may have driven those increments despite the changes both to cyclic backbones and *de novo* sequences. However, none of the CSP/cycRT39_3-conformers reached the highest ∼ 10^-8^ picoMolar affinities of CSP/RT39_3 (**Figure 3, open red circle**).

**Figure 3.**
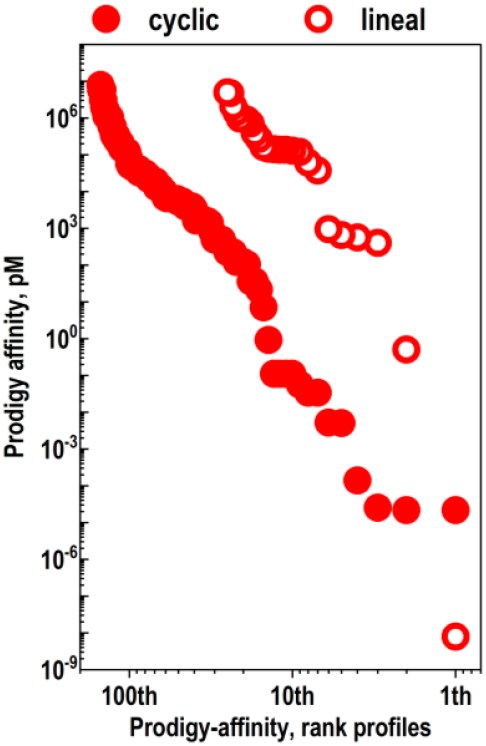
Affinity rank profiles of CSP/ALP- and CSP/cycALP-conformers. *Plasmodium falciparum* CSP (XM_001351086) 1-397 amino acid residue sequences were Alphafold2 co-modeled with the RT39, RT39M, RT53, RK37 and RK16 ALP sequences (**Table 1**). Five (_1 to _5) CSP/ALP conformers per ALP were generated. Cyclizations were performed by batch-modified af_ cyc_design.ipynb ^44^, to generate six (_0 to _5) cyclic-conformers per CSP/ALP. In both cases, the picoMolar (pM) affinities of the *de novo* generated models were approximated by Prodigy. For each lineal or cyclic conformer groups their affinities were merged to draw the overall rank profiles. Except the T39_3 conformer, all the CSP/ALP conformers increased their affinities after cyclization to CSP/cycALP conformers.

Each of the six CSP/cycRT39_3-conformers predicted affinities higher than any of those from the mutant CSP/cycRT39M-conformers (**Figure 4, RT39** and **RT39M**). These differences may suggest experimental membranolytic-killing and membranolytic-blocking activity correlations, respectively^**5**^. On the other hand, the limited affinities of both CSP/RT53- and their CSP/cycRT53-conformers may suggest alternative mechanisms for RT53 experimental partial membranolytic-killing activities^**5**^. Most CSP/cycRK16- and cycRK37-conformers predicted lower affinities than those of the CSP/cycRT39_3, suggesting those non-membranolytic ALP may have alternative weaker interactions with CSP.

**Figure 4.**
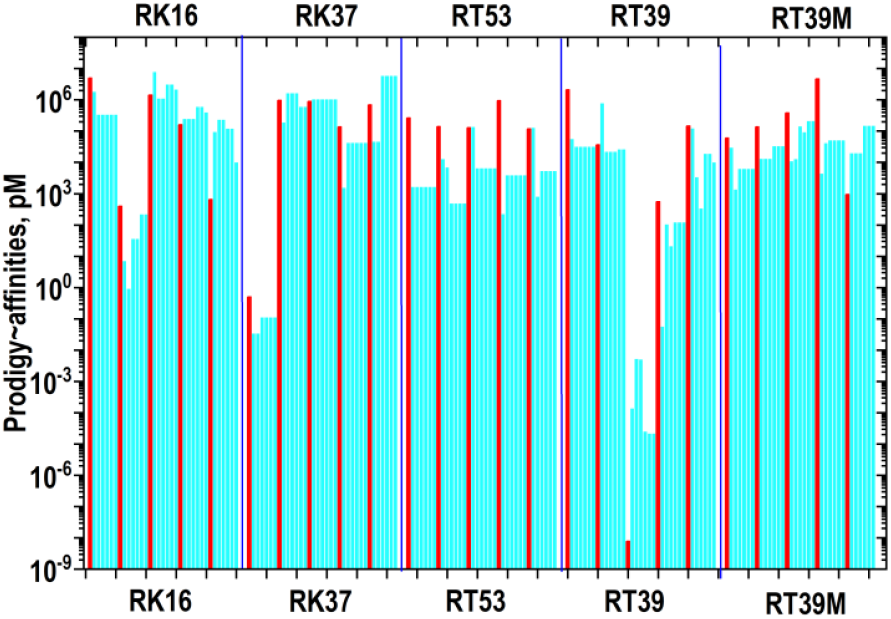
CSP/ALP (red) and their corresponding CSP/cycALP-conformer (cyan) affinities. *Plasmodium falciparum* full-length CSP (XM_001351086) 1-397 residues were co-modeled with ALP sequences (RK16, RK37, RT53, RT39, RT39M) (**Table 1, Figure 3**). The resulting CSP/ALP were cyclized using batch-modified af_ cyc_design. ipynb ^44^, to generate _0 to _5 CSP/cycALP-conformers per each ALP. Their affinities were approximated by Prodigy in PicoMolar (pM) using 4 Å distance contact predictions. **Red bars**, Five CSP/ALP-conformers per ALP. The conformer names were ordered from left to right (_1 to _5) **Cyan bars**, Six CSP/cycALP-conformers per CSP/ALP-conformers. The cyclic-conformer names were ordered from left to right (_0 to _5).

### Visualization of CSP/ALP conformers

PyMol visualization of the CSP/RT39 conformer complexes confirmed similar targeted tridimensional locations on CSP between ALP- and their cycALP-conformers, despite their different backbones, amino acid sequences and affinities (**Supplementary Materials / CSP+cyclic-pse.zip**). A comparative example between CSP/RT39_3 and one of its corresponding top CSP/cycRT39_3_3-conformer, illustrates that their small 3D differences were mostly at the bended terminal amino acids (**Figure 5 red cartoon**). Thus, the N-terminal ∼ 9 and the C-terminal ∼7 amino acids of the CSP/RT39_3 α-helix sequence (**Figure 5, green cartoon**), were bended to interconnect each other after cyclization in the CPS/cycRT39_3_3 conformer (**Figure 5, red cartoon**).

**Figure 5.**
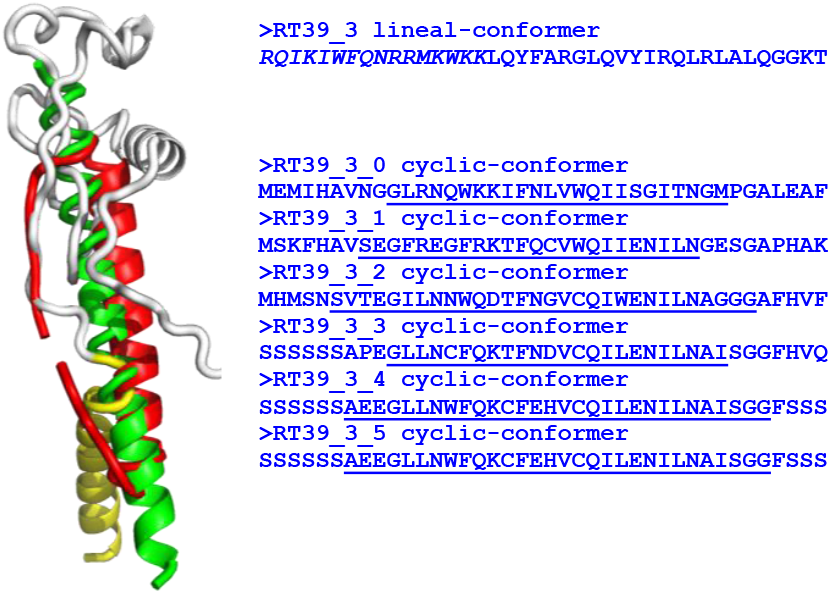
CSP/RT39_3 and CSP/cycRT39_3_3 conformers in complex with CSP-TSR. The CSP/RT39_3 conformer was merged with the CSP/cycRT39_3_3 ^44^ . Here only the CSP drawn was restricted to its TSR domain, the rest were hidden for clarity. All the CSP/ALP- and CSP/cycALP-conformers are included at **Supplementary Materials / CSP+lineal-pse.zip and CSP+cyclic-pse.zip**. The CSP/RT39_3_4- and _5-conformers predicted identical peptide amino acid sequences. **Yellow cartoon**, CSP-C-terminal α-helix (376-396 residues) **Gray cartoon**, CSP-TSR domain (312-324 residues) **Green cartoon**, RT39_3 α-helix ALP **Red cartoon**, RT39_3_3 cyclic-conformer (drawn open in PyMol cartoons, remains closed in PyMol sticks) **Right sequences**, fasta amino acid sequences saved from the corresponding pdb PyMol files. **Underlined sequences**,∼ residues of the cyclic-conformer sequences maintained as α-helices **Italic sequence**, RK16 penetratin at the RT39_3 lineal-conformer

The visual comparison of the CSP/RT39_3-conformer confirmed the coverage of the most important CSP domains (**Figure 1**), including the amino α-helix implicated in hepatocyte-binding, NANP disordered-repeats and the C-terminal TSR (**Supplementary Materials / CSP+cyclic-pse.zip**).

Due to the high affinities of the top CSP/cycRT39_3-conformers, it may be also possible to interact with CSP when in the mosquitos, shortly after CSP would be transferred to the circumsporozoite membranes. Would circumsporozoite targeting on mosquitos hemolymph / salivary glands with cycRT39 be possible?.

### Preliminary exploration of Uniprot <u>A</u>nti<u>m</u>icrobial <u>p</u>eptides

The chemical similarity between ALP and non-tested <u>**a**</u>nti<u>**m**</u>icrobial <u>**p**</u>eptide sequences (AMPs) for inhibition of *Plasmodium* circumsporozoites, prompted to preliminary exploration of their possible CSP docking. The number of AMPs included into Uniprot were of 82986 antimicrobial sequences (as accessing at September 2025). Therefore, a complete computational generation of CSP complex conformers, would require both large computational storage capacity and long computational times. On the other hand, because of the 39-mer optimal size for membranolytic-killing, excesive AMP sizes may not be of particular interest in the context of possible circumsporozoite inhibitors.Therefore, Alphafold2 CSP/AMP co-models were generated in selected AMP ranges from 30-to 65-mer. Because of our actual computational restrictions, batches of ∼ 300 AMPs per batch were used to generate a total of 1818 CSP/AMP conformers (one per AMP). The resulting CSP/AMP-conformers predicting < 1 pM affinities resulted in 28 top CSP/AMP-conformers, corresponding to AMPs between 47-to 51-mer (**Figure 6A**) (**Supplementary materials / 28AMPtops-pse.zip**).

**Figure 6.**
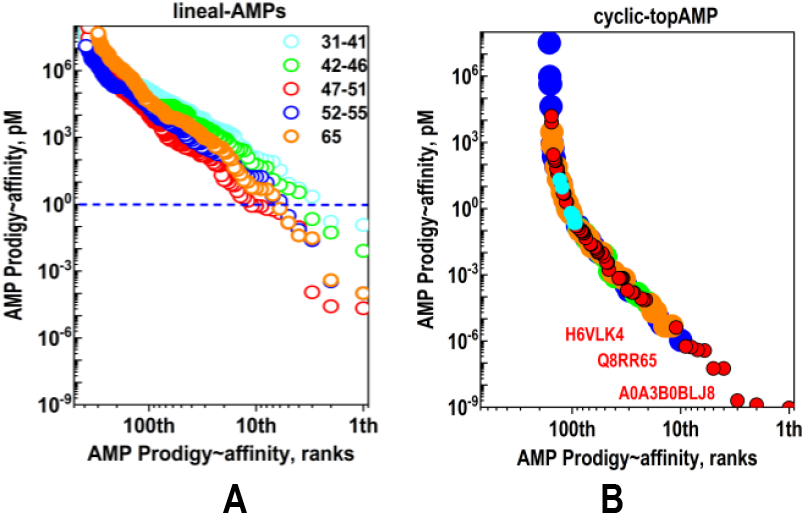
Affinity rank profiles of CSP/lineal- and top cyclic-conformers from Uniprot AMPs. **A) CSP/lineal-AMPS**. Alphafold2 co-modeling was performed with CSP and AMP lineal-peptide sequences downloaded from Uniprot by selecting different size peptides (number of amino acids indicated in the insert as open circle colors: **cyan, green, red, blue** and **orange**). **Blue dashed line**. Threshold affinities < 1 pM served to define 28 top CSP/AMPs co-modeled complexes with expected higher probabilities of improvement after cyclization (**Supplementary Materials / 28AMPtops-pse.zip**). **B) CSP/top cyclic-AMPs**. Cyclization of the top CSP/AMPs generated six conformer/sequences per top lineal-AMPs merged for the rank profiles. The colours correspond to sizes at the insert of **Figure 6A**. The highest affinities corresponded to five conformers/sequences derived from **A0A3B0BLJ8**.

None of the CSP/AMP-conformers predicted higher affinities than CSP/RT39). However, cyclization of some of the top CSP/AMPs predicted conformer affinities similar to those of CSP/RT39 (**Figure 6B**). The highest affinities corresponded to five of the six CSP/cycA0A3B0BLJ8-conformers. The AMP A0A3B0BLJ8 corresponds to *Lantibiotic*, a 49-mer peptide from *Streptococcus chosunensis*, which induces membranolytic-killing of Gram-positive bacteria. Other of the highest affinity conformers corresponded to CSP/cycH6VLK4, another *Lantibiotic* H6VLK4 AMP of 49-mer from *Streptococcus salivarius*, inducing membrane pores in Gram-positive bacteria, and to CSP/cycQ8RR65, derived from an AMP Q8RR65 *bactericine* of 50-mer from *Enterococcus mundtii*.

These preliminary AMP screening results suggests that the ∼ 49-mer may be the optimal size to find other still unexplored CSP/cyclic-conformers among the abundant Uniprot AMPs. On the other hand, a more accurate and wide screening of the remaining AMP sequences and their cyclic-conformers may deserve further screenings once computational methods would be refined.

## Computational limitations

The main reasons to explore for peptide cyclizations were to reduce degradation by exoproteases and to increase chemical stability. However, improvements on affinities were also predicted. Although high in the protein-protein complexes used for training (∼ 0.7 correlations, similar to experimental data), the expected Prodigy predictions, like those from any other affinity calculation program are less accurate when applied to unrelated protein-protein complexes, like those generated during this work. Nevertheless, comparison between relative rather than absolute predictions may still be valid because all complexes contained ∼ 90 % of identical CSP amino acid sequences. Estimations of absolute protein-protein affinities would always remain unaccurate, unless more numerous and representative biological activities could be used to train deep-learning approaches. Additionally, biological activities may be also influenced by the simplifications included by the selected computational constrains, such as the absence of water interactions, CSP model dependence, possible resistant mutations, limited CSP-domain targetings and/or other yet unknowns. Despite all mentioned limitations, the computationally top cyclic-conformers newly described here and/or others yet to be discovered among the enormous unexplored peptide space^36, 39^, may offer alternative candidates for *in vitro* and *in vivo* validations.

## Conclusions

The *P*.*falciparum* circumsporozoite membranolytic-killing RT39 ALP described before has demonstrated higher computational affinity to CSP for 20% of the Alphafold2 generated conformers. No other previously described ALPs reached such CSP high affinities. These findings suggest that docking to CSP could contribute to explain the RT39 circumsporozoite high membranolytic-killing activity. Additionally, most computational CSP/cycRT39 and some of CSP/cycAMPs newly identified by preliminary screening of AMPs, predicted high affinities to CSP. For practical applications, cyclic-peptides targeting the external circumsporozoite CSP are easier-to-synthesize, proteolytic more resistant, chemically more stable and cell permeability-independent^43-50^, compared to other lineal-peptides and other small drug-like alternatives. The combination of these advantages could made cyclic-peptides good candidates to target circumsporozoite during their migrations whether inside mosquitos and/or humans.

## Computational methods

### *Plasmodium falciparum* circumsporozoite CSP Alphafold2 co-models

The **c**ircum**s**porozoite **p**rotein (CSP) XM_001351086 sequence of the *P*.*falciparum* 3D7 including its crystallized C-terminal 301-396 residues (3vdj.pdb)^45^ were Alphafold2 co-modeled ^45,4, 26, 47^ with the sequences of the previously described **a**mphipathic **l**ineal-**p**eptides (ALP) RT39, RT39M, RT53, RK37 and RK16 tested for circumsporozoite membranolytic-killing (**Table 1**)^5^ to generate CSP/ALP-conformers. The Alphafold2 ipynb code was slighly modified to include Driver mount at the first cell and automatically Driver downloaded-saved the zipped results at the final cell (**Supplementary Materials / AlphaFold2batchpdb4(5)ipynb.zip**). Once the results.zip were transferred into the local computer environment, a home-designed PyMol/Python script unzipped the generated CSP/ALP pdb files for Prodigy affinity estimations.

Alphafold2 CSP/AMP co-models were similarly constructed by downloading Uniprot **a**nti**m**icrobial **p**eptides (AMPs) in ranges from 30-to 65-mer amino acids conserving their Uniprot ID names. A Geminis-aided Python script was developed to automatically select, download the Uniprot peptides desired sizes or size ranges. The corresponding CSP/AMP-conformers were generated from the amino acid sequences in one letter-code to be submitted in batches of ∼ 300 AMPs to Alphafold2 co-modelling (**Supplementary Materials / AlphaFold2batchpdb4(5)ipynb.zip**). Result.zip files were analysed as above.

### Estimation of CSP/peptide complex affinities and amino acid contacts by Prodigy

Validation of the Alphafold2-generated CSP/ALP- and CSP/AMP-conformer pdbs, calculation of their interatomic interface distances, and buried surface area, were calculated by Prodigy^56-60^ using Biopython https://www.biopython.org), freesasa (https://github.com/mittinatten/freesasa), and naccess (http://www.ncbi.nlm.nih.gov/pubmed/994183), respectively. A local Prodigy anaconda3-guided Python 3.10.8 installation on Windows 10 was run to calculate short-interactions using “-- distance cutoff” of 4 Å^42^. Using batches as described^34,39.^ For each complex-conformer outputs included: all_pdb/, all_contacts/ (text file), all_pml (to draw side chains in PyMol), all_Kcal (docking affinities in Kcal/mol), orderpdb (pdb ordered by their Kcal per mol) and all_kcalpM.txt including calculated picoMolar affinities (10^12^*^(exp(Kcal/0.592).

### Lineal-peptide cyclization

Head-to-tail cyclization of the lineal-peptides (N-C terminal bonds) were generated in batch using a modified Colab notebook developed from the af_ cyc_design.ipynb ^44^ https://github.com/sokrypton/ColabDesign/blob/main/af/examples/af_cyc_design.ipynb. The notebook was provided with additional batch fixbb protocol details as most recently described ^4^. Results were automatically downloaded-saved at Drive and their ^*^.pdbs submitted to Prodigy and PyMol for analysis.

## Supporting information

28AMPtops-pse

AlphaFold2batchpdb4(5)ipynb

CSP+cyclic-pse

CSP+lineal-pse

GraphycalAbstract-pse

## Supplementary information

**Table S1.**
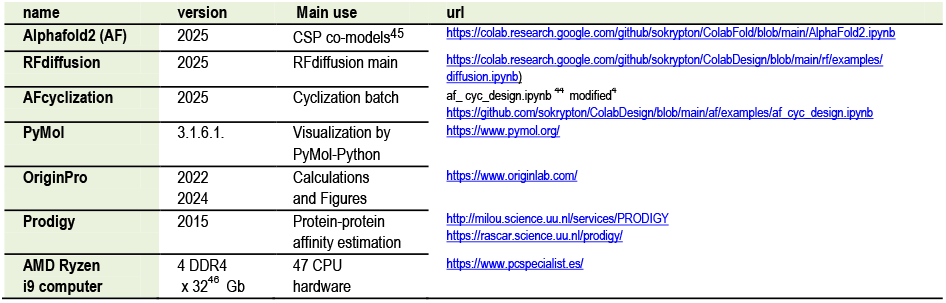
Computational software.

## Supplementary Materials

### GraphycalAbstract-pse.zip

The full-length mRNA of *Plasmodium falciparum* CSP was co-modeled by Alphafold2 with the RT39 lineal- and cyclic-peptides to generate docking conformers. **Gray cartoons**, predicted CSP 3D partial structure (most CSP NANP-disordered repeats were eliminated for clarity). **Yellow α-helix**, CSP C-terminal transmembrane TM α-helix. **Green cartoon**, RT39 lineal-peptide conformer. **Red cartoon**, head-to-tail cyclic-conformer derived from RT39 in complex with CSP (PyMol stick representation). Cyclization generated both cyclic backbones and *de novo* amino acid sequences.

### CSP+lineal-pse.zip

The file contains zipped Alphafold2-generated five conformers for each of the ALPs (RT39, RT39M, RT53, RK37 and RK16) complexed with CSP. To best visualization of the conformer ALPs, CSP were drawn in **gray** while ALP conformers were drawn in **green**. The ^*^.pse files can be opened in the latest versions of PyMol 3.

### CSP+cyclic-pse.zip

The file contains zipped Alphafold2-generated six cyclic conformers from ALPs and their *de novo* sequences for each ALP complexed with CSP (CSP1+RT39.pse, CSP1+RT39M.pse, CSP1+RT53.pse, CSP1+RK37.pse and CSP1+RK16.pse). To best visualization of the cyclic ALP conformers, they could be re-drawn in **red** and CSP could be re-drawn in **gray** similarly to CSP1+RT39 and CSP1+RT39M. The ^*^.pse files can be opened in the latest versions of PyMol 3.

### 28AMPtops-pse.zip

Uniprot antimicrobial AMP sequences in the 30-65-mer sizes were Alphafold2 complexed with CSP in batches of ∼ 300 AMPs per batch (1818 AMP total number of sequences). The resulting CSP/AMP co-models predicted maximal numbers of conformers with < 1 pM affinity for 28 top CSP/AMPs corresponding to AMP sizes between 47-to 65-mer. None of the CSP/AMPs conformers predicted higher affinities than RT39. Most of their CSP amino acid contacts targeted their TSR domains. All the complexes were aligned to their TM transmembrane 376-396 sequences (α-helix in **yellow**) in PyMol using the commands: “sele TM,A0A3B0BLJ8 and resi 376-396” and then “TM/A/align/all to this (^*^/CA)”.

### AlphaFold2batchpdb4(5)ipynb.zip

This is a Colab notebook ipynb modified for this work to perform batches of ∼300 lineal-AMP peptides, automatically saving results as zipped pdb-only files to Drive and after-run-automatically cleaning of 5-7 Gb of files which were generated after running/finishing each batch (one practical problem for large numbers of folds). After modelling, all the lineal-peptide ^*^.pdbs were submitted to Prodigy and PyMol for further analysis of affinities and contacts.

## Authors’ contributions

JC designed, performed and analyzed the computational work and drafted the manuscript.

## Acknowledgements

Thanks are due to the participating people at the Slack-Boltz forum (https://boltz-community.slack.com/ssb/redirect) and the Alphafold Discord forum (https://discord.com/channels/).

