## Supplementary material for "*Plasmodium* killing and antimicrobial lineal or cyclic-peptides computationally targeted the circumsporozoite CSP protein": AlphaFold2batchpdb4(5)ipynb: AlphaFold2batchpdb4(5).ipynb

```
# %% [markdown]
#AlphaFoldbatchpdb4(5).ipynb ColabFold v1.5.5: AlphaFold2 w/ MMseqs2 BATCH
<img src="https://raw.githubusercontent.com/sokrypton/ColabFold/main/.github/ColabFold_Marv_Logo_Small.png" height="100" align="right" style="height:256px">
[Mirdita M, Schütze K, Moriwaki Y, Heo L, Ovchinnikov S, Steinegger M. ColabFold: Making protein folding accessible to all.
*Nature Methods*, 2022](https://www.nature.com/articles/s41592-022-01488-1)
modified to extract pdb batches information only by J.Coll 4/09/2025

# %% [code]
#@title 1. MOUNT GOOGLE DRIVE
from google.colab import drive
drive.mount('/content/drive')
from sys import version_info
python_version = f"{version_info.major}.{version_info.minor}"

# %% [code]
#@title 2. INPUTS
input_dir = '/content/drive/MyDrive/AMP/fasta65' #@param {type:"string"}
result_dir = '/content/drive/MyDrive/AMP/result299' #@param {type:"string"}

#@markdown ---
#@markdown ### Advanced settings
msa_mode = "MMseqs2 (UniRef+Environmental)" #@param ["MMseqs2 (UniRef+Environmental)", "MMseqs2 (UniRef only)","single_sequence","custom"]
num_models = 1 #@param [1,2,3,4,5] {type:"raw"}
num_recycles = 1 #@param [1,2,3,6,12,24,48] {type:"raw"}
stop_at_score = float("30") #@param {type:"string"}
##@markdown - early stop computing models once score > threshold (avg. plddt for "structures" and ptmscore for "complexes")
use_custom_msa = False
num_relax = 0 #@param [0, 1, 5] {type:"raw"}
use_amber = num_relax > 0
relax_max_iterations = 100 #@param [0,100,200,2000] {type:"raw"}
use_templates = False #@param {type:"boolean"}
do_not_overwrite_results = True #@param {type:"boolean"}
zip_results = True #@param {type:"boolean"}
print("input_dir: ", input_dir)
print("result_dir: ", result_dir)
print("msa_mode: ", msa_mode)
print("num_models: ", num_models)
print("num_recycles: ", num_recycles)
print("stop_at_score: ", stop_at_score)


# %% [code]
#@title 3. INSTALL DEPENDENCIES
%%bash -s $use_amber $use_templates $python_version
set -e
USE_AMBER=$1
USE_TEMPLATES=$2
PYTHON_VERSION=$3
if [ ! -f COLABFOLD_READY ]; then
  # install dependencies
  # We have to use "--no-warn-conflicts" because colab already has a lot preinstalled with requirements different to ours
  pip install -q --no-warn-conflicts "colabfold[alphafold-minus-jax] @ git+https://github.com/sokrypton/ColabFold"
  if [ -n "${TPU_NAME}" ]; then
    pip install -q --no-warn-conflicts -U dm-haiku==0.0.10 jax==0.3.25
  fi
  ln -s /usr/local/lib/python3.*/dist-packages/colabfold colabfold
  ln -s /usr/local/lib/python3.*/dist-packages/alphafold alphafold
  # hack to fix TF crash
  rm -f /usr/local/lib/python3.*/dist-packages/tensorflow/core/kernels/libtfkernel_sobol_op.so
  touch COLABFOLD_READY
fi
# Install appdirs
pip install -q appdirs #erroneously added when input was 0 or required?
# Download params (~1min)
python -m colabfold.download
# setup conda
if [ ${USE_AMBER} == "False" ] || [ ${USE_TEMPLATES} == "False" ]; then #not needed for batch
  if [ ! -f CONDA_READY ]; then
    wget -qnc https://github.com/conda-forge/miniforge/releases/download/25.3.1-0/Miniforge3-25.3.1-0-Linux-x86_64.sh
    bash Miniforge3-25.3.1-0-Linux-x86_64.sh -bfp /usr/local 2>&1 1>/dev/null
    rm Miniforge3-25.3.1-0-Linux-x86_64.sh
    conda config --set auto_update_conda false
    touch CONDA_READY
  fi
fi
# setup template search
if [ ${USE_TEMPLATES} == "True" ] && [ ! -f HH_READY ]; then
  conda install -y -q -c conda-forge -c bioconda kalign2=2.04 hhsuite=3.3.0 python="${PYTHON_VERSION}" 2>&1 1>/dev/null
  touch HH_READY
fi
# setup openmm for amber refinement
if [ ${USE_AMBER} == "True" ] && [ ! -f AMBER_READY ]; then
  conda install -y -q -c conda-forge openmm=8.2.0 python="${PYTHON_VERSION}" pdbfixer 2>&1 1>/dev/null
  touch AMBER_READY
fi

#import os
#from colabfold.batch import get_queries
#print(f"Current input_dir: {input_dir}")
#print(f"\nListing contents of {input_dir}:")
#!ls -l "{input_dir}"
#print("\nOutput of get_queries:")
#queries, is_complex = get_queries(input_dir)
#print(f"Queries found: {queries}")
#print(f"Is complex: {is_complex}")

# %% [code]
#@title 4. RUN PREDICTION
import sys
from colabfold.batch import get_queries, run
from colabfold.download import default_data_dir
from colabfold.utils import setup_logging
from pathlib import Path
# For some reason we need that to get pdbfixer to import
if use_amber and f"/usr/local/lib/python{python_version}/site-packages/" not in sys.path:
    sys.path.insert(0, f"/usr/local/lib/python{python_version}/site-packages/")
setup_logging(Path(result_dir).joinpath("log.txt"))
queries, is_complex = get_queries(input_dir)
run(
    queries=queries,
    result_dir=result_dir,
    use_templates=use_templates,
    num_relax=num_relax,
    relax_max_iterations=relax_max_iterations,
    msa_mode=msa_mode,
    model_type="auto",
    num_models=num_models,
    num_recycles=num_recycles,
    model_order=[1, 2, 3, 4, 5],
    is_complex=is_complex,
    data_dir=default_data_dir,
    keep_existing_results=do_not_overwrite_results,
    rank_by="auto",
    pair_mode="unpaired+paired",
    stop_at_score=stop_at_score,
    zip_results=zip_results,
    user_agent="colabfold/google-colab-batch",
)

# %% [code]
#@title 5. EXTRACT PDBs, ZIP & DRIVE SAVE
import os
import zipfile
import shutil
import tempfile

# Construct the full path to the result_dir
result_dir_path = result_dir
all_files = os.listdir(result_dir_path)
zip_files = [f for f in all_files if f.endswith('.result.zip')]

# Create a temporary directory to extract PDBs to
with tempfile.TemporaryDirectory() as tmpdir:
    print(f"Using temporary directory: {tmpdir}")

    # Extract only PDB files from each zip into the temporary directory
    for zip_filename in zip_files:
        zip_filepath = os.path.join(result_dir_path, zip_filename)
        if os.path.exists(zip_filepath):
            with zipfile.ZipFile(zip_filepath, 'r') as zip_ref:
                for file_in_zip in zip_ref.namelist():
                    if file_in_zip.endswith('.pdb'):
                        # Construct the full path for extraction in the temp directory
                        extracted_path = os.path.join(tmpdir, os.path.basename(file_in_zip))
                        # Extract the file to the temporary directory
                        with open(extracted_path, 'wb') as outfile:
                            with zip_ref.open(file_in_zip) as infile:
                                outfile.write(infile.read())
            print(f"Extracted PDBs from '{zip_filename}' to temporary directory.")
        else:
            print(f"Warning: Zip file not found at '{zip_filepath}'")

    # List files extracted to the temporary directory (for verification)
    extracted_pdb_files = [f for f in os.listdir(tmpdir) if f.endswith('.pdb')]
    print(f"\nFound {len(extracted_pdb_files)} PDB files in the temporary directory:")
    # for pdb_file in extracted_pdb_files:
    #     print(pdb_file) # Optional: print all filenames

    # Remove the common suffix from the filenames
    common_suffix = "_unrelaxed_rank_001_alphafold2_multimer_v3_model_1_seed_000"
    renamed_pdb_files = []
    for pdb_filename in extracted_pdb_files:
        if pdb_filename.endswith(common_suffix + '.pdb'):
            new_filename = pdb_filename.replace(common_suffix, "")
            original_filepath = os.path.join(tmpdir, pdb_filename)
            new_filepath = os.path.join(tmpdir, new_filename)
            os.rename(original_filepath, new_filepath)
            renamed_pdb_files.append(new_filename)
        else:
            renamed_pdb_files.append(pdb_filename) # Keep files that don't match the suffix
    print(f"All PDB files ending with '{common_suffix}' have been renamed.")


    # Define the name for the new zip file
    output_zip_filename = os.path.basename(result_dir_path) + '_all_pdbs.zip'
    output_zip_path_temp = os.path.join(tmpdir, output_zip_filename)

    # Create a new zip file containing all extracted PDBs
    print(f"\nCreating new zip file: '{output_zip_filename}'")
    with zipfile.ZipFile(output_zip_path_temp, 'w') as new_zip:
        for pdb_file in renamed_pdb_files: # Use the list of renamed files
            pdb_filepath_temp = os.path.join(tmpdir, pdb_file)
            new_zip.write(pdb_filepath_temp, arcname=pdb_file) # Add file to zip with just its name
    print(f"New zip file created at '{output_zip_path_temp}'")

    # Define the destination path in Google Drive
    destination_path = '/content/drive/MyDrive/AMP/' + output_zip_filename

    # Move the new zip file to Google Drive
    print(f"Moving '{output_zip_path_temp}' to '{destination_path}'")
    shutil.move(output_zip_path_temp, destination_path)
    print(f"Moved '{output_zip_filename}' to Google Drive.")

# The temporary directory and its contents are automatically removed after exiting the 'with' block
print("\nTemporary directory cleaned up.")

# Optional: List the contents of the result_dir to show original zips remain
# print(f"\nContents of '{result_dir_path}' after processing:")
# contents = os.listdir(result_dir_path)
# for item in contents:
#     print(item)

# %% [code]
#@title 6. CONFIRM PDBs DOWNLOAD & CLEAN UP RESULT_DIR
import os
import shutil

# Define the path to the final zipped PDB file in Google Drive
output_zip_filename = os.path.basename(result_dir) + '_all_pdbs.zip'
destination_path = '/content/drive/MyDrive/AMP/' + output_zip_filename

# Check if the final zip file exists in the destination
if os.path.exists(destination_path):
    print(f"Final zip file found at '{destination_path}'. Proceeding with cleanup.")

    # Clean up the contents of the result_dir
    print(f"Cleaning up contents of '{result_dir}'...")
    try:
        # Get list of items in result_dir
        items_to_delete = os.listdir(result_dir)
        if items_to_delete:
            for item in items_to_delete:
                item_path = os.path.join(result_dir, item)
                if os.path.isfile(item_path):
                    os.remove(item_path)
                    # print(f"Deleted file: {item}") # Uncomment for verbose deletion
                elif os.path.isdir(item_path):
                    shutil.rmtree(item_path)
                    # print(f"Deleted directory: {item}") # Uncomment for verbose deletion
            print(f"Contents of '{result_dir}' cleaned up.")
        else:
            print(f"'{result_dir}' is already empty.")
    except Exception as e:
        print(f"An error occurred during cleanup: {e}")

else:
    print(f"Final zip file not found at '{destination_path}'. Skipping cleanup of '{result_dir}'.")
    print("Please ensure the previous step successfully created and moved the zip file.")
```
